# Connectivity dynamics from wakefulness to sleep

**DOI:** 10.1101/380741

**Authors:** Eswar Damaraju, Enzo Tagliazucchi, Helmut Laufs, Vince D Calhoun

## Abstract

Interest in time-resolved connectivity in fMRI has grown rapidly in recent years. The most widely used technique for studying connectivity changes over time utilizes a sliding windows approach. There has been some debate about the utility of shorter versus longer windows, the use of fixed versus adaptive windows, as well as whether observed resting state dynamics during wakefulness may be predominantly due to changes in sleep state and subject head motion. In this work we use an independent component analysis (ICA)-based pipeline applied to concurrent EEG/fMRI data collected during wakefulness and various sleep stages and show: 1) connectivity states obtained from clustering sliding windowed correlations of resting state functional network time courses well classify the sleep states obtained from EEG data, 2) using shorter sliding windows instead of longer non-overlapping windows improves the ability to capture transition dynamics even at windows as short as 30 seconds, 3) motion appears to be mostly associated with one of the states rather than spread across all of them 4) a fixed tapered sliding window approach outperforms an adaptive dynamic conditional correlation approach, and 5) consistent with prior EEG/fMRI work, we identify evidence of multiple states within the wakeful condition which are able to be classified with high accuracy. Classification of wakeful only states suggest the presence of time-varying changes in connectivity in fMRI data beyond sleep state or motion. Results also inform about advantageous technical choices, and the identification of different clusters within wakefulness that are separable suggest further studies in this direction.

## 1. Introduction

The human brain continuously engages in mental activities that include introspection, theory of mind, and future planning, even when not actively pursuing a task. Fluctuations during the resting state have been shown to show functional network topology similar to that observed in task imaging studies (Calhoun et al., 2008; Smith et al., 2009). Since the first reports on time varying functional connectivity during task performance (Sakoǵlu et al., 2010) and during a typical resting functional magnetic resonance imaging (fMRI) scan (Chang and Glover, 2010; Sakoǵlu et al., 2010), there has been interest in studying and characterizing the changes in functional connectivity between brain regions on a shorter time scale, also referred to as time-resolved or dynamic functional connectivity(Calhoun et al., 2014; Hutchison et al., 2013). Recent years have seen a sharp increase in development of novel methods to characterize these dynamics (Allen et al., 2012a; Cribben et al., 2013; Kang et al., 2011; Tagliazucchi et al., 2012a; Vidaurre et al., 2017; Yaesoubi et al., 2015a)(see (Preti et al., 2017) for a thorough review).

One popular method to estimate connectivity dynamics in functional neuroimaging data is the use of sliding windows to estimate the connectivity between regions/networks. These time-varying connectivity estimates, when computed on the time courses from brain regions or seeds defined a priori are referred to as dynamic functional connectivity (dFC) and those estimated from the time courses of networks obtained from blind decomposition techniques such as independent components analysis (ICA) are referred to as dynamic functional network connectivity (dFNC). In this work we focus on the latter, which leverages the benefits of ICA including data-driven identification of networks, robustness to noise, separation of overlapping signals of interest, and estimation of homogeneous networks that capture individual subject variability (Calhoun and de Lacy, 2017; Yu et al., 2017). Using an ICA based pipeline, Allen et al. (2012a) reported the presence of stable, recurring connectivity patterns/states obtained from clustering pairwise correlation estimates of windowed time courses of ICA derived intrinsic networks from resting fMRI data and more recently (Abrol et al., 2017) showed connectivity patterns and dynamic metrics are highly replicable across multiple independent data sets. However, a few studies have raised concerns over the quality of connectivity estimates obtained using the sliding window method (Lindquist et al., 2014; Smith et al., 2011), choice of window size (Hindriks et al., 2016; Leonardi and Van De Ville, 2015; Sakoǵlu et al., 2010; Zalesky and Breakspear, 2015), and the ability of the method to capture meaningful state transitions (Shakil et al., 2016). Others have primarily attributed the observable changes in connectivity in resting fMRI data to sleep state, sampling variability and head motion (Laumann et al., 2016).

In our prior concurrent EEG/fMRI work (Allen et al., 2017), windows corresponding to distinct dFNC connectivity states estimated using a sliding-window correlation method were associated with distinct electrophysiological signatures during both eyes open and eyes closed awake conditions and showed the ability of the method to track subject vigilance. However, since subjects were mostly awake throughout the scan sessions, simultaneously acquired electroencephalogram (EEG) data did not show enough state transitions from wakefulness to assess how well observed dFNC state transitions correspond to neurobiological state transitions i.e. observable changes in subject sleep state from EEG data.

Here we use simultaneous EEG-resting fMRI data collected continuously over 50 minutes while the subjects transitioned between wakefulness and different sleep stages (defined by EEG-based sleep scoring) and assessed the ability of sliding window based dFNC measures to track the changes across these different wakefulness states. In addition to comparing the dFNC measure to sleep, this study provides us an opportunity to evaluate the impact of several technical choices within a real world dataset rather than in simulations (Leonardi and Van De Ville, 2015; Sakoǵlu et al., 2010). For example, we compare the impact of the length of the sliding window on our ability to predict sleep stage. Windows should be short enough to be a good compromise between the ability to capture time varying connectivity without being too sensitive to noise. In addition, we compare a ‘fixed length’ sliding window approach to a method from econometrics which has been applied to fMRI data, the dynamic conditional correlation (DCC) (Engle, 2002; Lindquist et al., 2014). DCC uses an adaptive window size and has been reported to show better test-rest reliability compared to ‘fixed length’ sliding-window methods in estimating time varying functional connectivity (Choe et al., 2017). We also evaluate the relationship of the estimated states to motion, in particular we were interested in whether all states would show a similar relationship to motion or whether a subset of states captures the variance associated with motion.

## 2. Methods

### 2.1. Data acquisition

Resting state functional MRI data was collected from 55 subjects for 52 minutes each (1505 volumes of echo planar images, repetition time (TR)=2.08 s, TE = 30 ms, matrix 64 x 64; FOV = 192 mm^2^, Thickness=2 mm, 1 mm gap between slices) on a Siemens 3T Trio scanner while the subjects rested with their eyes closed (details in (Tagliazucchi et al., 2012b)). Simultaneous EEG was acquired on 30 EEG channels with FCz as the reference (sampling rate = 5 kHz, low pass filter = 250 Hz, high pass filter = 0.016 Hz) using an MR compatible device (BrainAmp MR+, BrainAmp ExG; Brain Products, Gilching, Germany).

### 2.2. Data preprocessing

EEG data underwent MRI and pulse artifact correction based on average artifact subtraction (Allen et al., 1998), followed by ICA-based residual artifact rejection. This data was subsequently sleep staged into wakeful (W), drowsy/light sleep (N1), moderate sleep (N2), and deep sleep (N3) stages by a sleep expert per American Academy of Sleep Medicine (AASM) criteria (AASM and Iber, 2007) resulting in a hypnogram for the scan duration for each subject. None of the participants went into REM sleep stage during the scan period. The first 5 volumes of functional imaging data were discarded to account for T1 equilibration effects. The data were then corrected for head movement (rigid body) and slice timing differences. Subject data was then spatially normalized to MNI template space using SPM12 toolbox and resampled to 3 mm^3^ isotropic voxel resolution. Since the scan duration was long, we detrended each voxel time series using a high model order (21) polynomial. The subject brain voxel data were then spatially smoothed to 5 mm FWHM using AFNI’s 3dBlurInMask. Finally, the voxel time courses were variance normalized (z-scored).

### 2.3. Independent component estimation

We performed a group independent component analysis (Calhoun et al., 2001; Calhoun and Adali, 2012) using the GIFT toolbox (http://mialab.mrn.org/software/gift) to decompose the data into 100 spatially independent components each associated with a coherent time course. First subject data were reduced in time to 1400 points using principal components analysis (PCA). These time reduced datasets were then concatenated in time and a group PCA was used to further reduce the data to 100 orthogonal directions of maximal variance. Spatial ICA analysis was performed on this data to obtain 100 spatially independent components using infomax algorithm. To ensure stability of decomposition, the Infomax ICA algorithm was repeated 20 times in ICASSO (Himberg et al., 2004) with random initialization, and aggregate spatial maps were estimated as the modes of the component clusters. Subject specific component maps and their associated time courses were obtained using spatio-temporal regression (Erhardt et al., 2011).

### 2.4. Component selection and dynamic FNC estimation

Of the 100 spatial ICA components, we identified 62 components as intrinsic connectivity networks (ICNs) in a semi-automatic manner using the spatial profile of component maps, and dominant low frequency spectral power of their time courses as described in (Allen et al., 2011). The corresponding ICN time courses of each subject were orthogonalized against estimated head movement parameters in a regression framework and then filtered using 5th order Butterworth filter with a passband of 0.01 to 0.15 Hz. Following (Allen et al., 2012a), we performed dynamic functional network connectivity analysis using the post processed ICN time courses. Briefly, we computed pairwise correlations between tapered windowed segments (a rectangular window of 30 TR (60s) convolved with a Gaussian kernel of *σ* = 3s) of time courses sliding in steps of 1 TR. This resulted in 1500-30=1470 windows over which we estimated dFNC between 62 independent network time courses (1891 pairs). Since the number of samples are smaller than a static approach using all the timecoure information, we used a robust estimation strategy employing the graphical LASSO method (Friedman et al., 2008) by placing a penalty on the L1 norm of the precision matrix (inverse correlation matrix) to promote sparsity. Given the concerns with estimation of dFNC using shorter windows (Leonardi and Van De Ville, 2015; Lindquist et al., 2014; Smith et al., 2011), we repeated the analysis with longer tapered window sizes of size 45 TR, 60 TR, and shorter tapered window sizes of size 16 TR and 22 TR.

### 2.5. K-means clustering and comparison to EEG hypnogram

The dFNC windows from a given sliding window were clustered using the K-means algorithm in two steps consistent with our previous work (Allen et al., 2012a). We first computed a time course of standard deviation of dFNC matrices for each subject and selected subset of subject windows corresponding to local maxima in standard deviation as subject exemplars. These subset exemplars were clustered using K-means algorithm with Manhattan (L1) distance as distance measure. The number of clusters was set to 5 based on elbow criterion of ratio of within to between cluster distance of windows from cluster centroids, referred to as dFNC states. The obtained centroids were used as starting points to cluster all the data. This two step procedure, one to identify initial starting points and subsequent clustering of all window data using these starting points results in stable centroid patterns across independent datasets. The K-means clustering (a hard clustering approach) assigns each dFNC window to one of the dFNC states. This assignment of subject windows in time to dFNC states results in a discrete vector referred to as state vector. Figure 1 shows the schematic of the analysis pipeline used in the paper. We assessed the correspondence between subject dFNC state vector and EEG-derived hypnogram using correlation. To further visualize the correspondence between dFNC estimates from the sliding-window method and EEG-based subject hypnogram, we project the multidimensional (1891) dFNC data into 2 dimensions using the t-distributed stochastic neighborhood embedding (t-SNE) algorithm that preserves the distances between similar objects in high dimensional space to low dimensions (van der Maaten et al., 2008). We also sought to characterize the temporal properties of state vectors obtained from each modality by computing state transition probabilities from these vectors and then compare the similarities between the two vectors.

**Figure 1:**
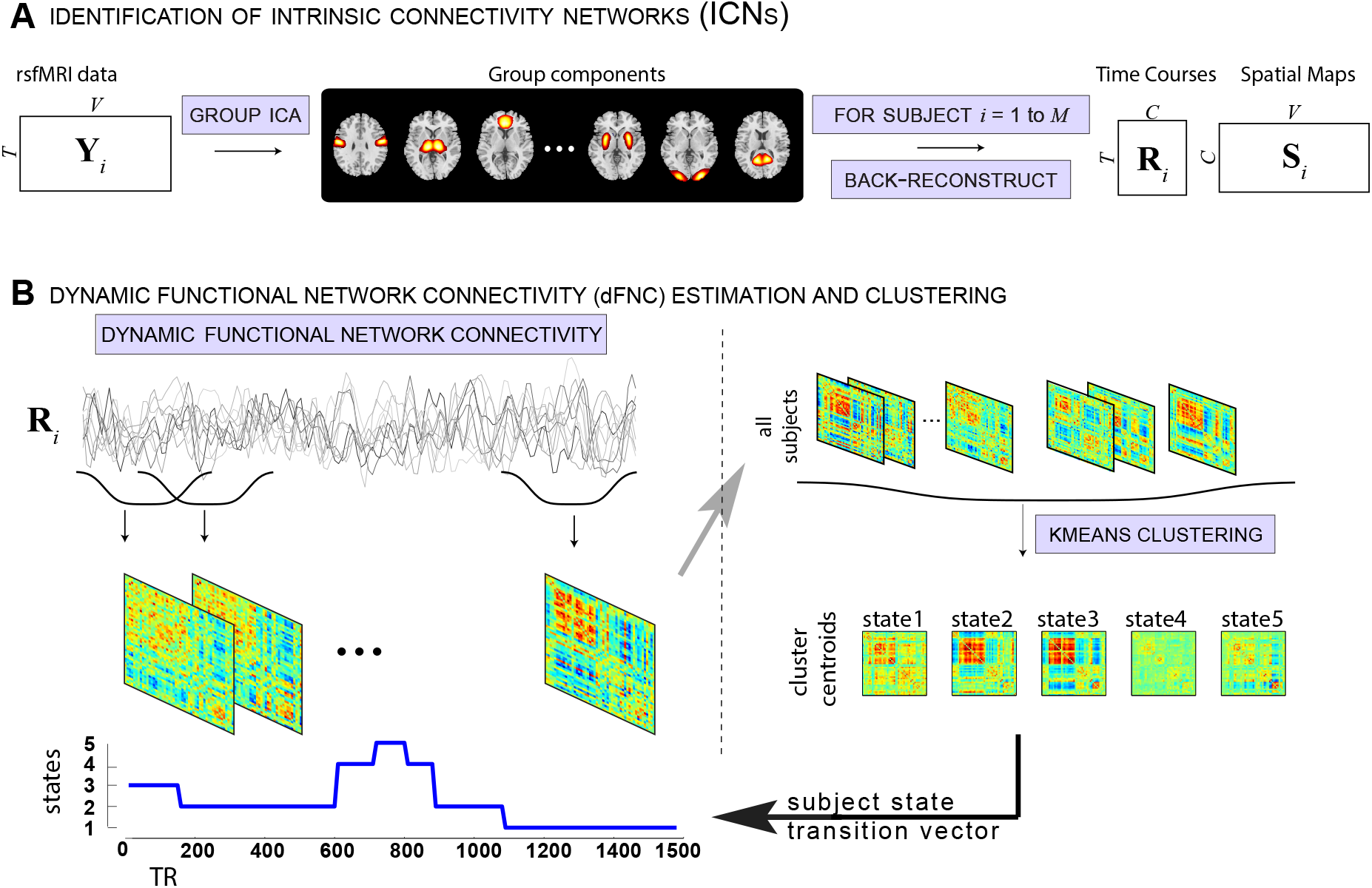
Schematic depicting resting fMRI data processing (adapted from (Damaraju et al., 2014)).

In our earlier work (Allen et al., 2017), we aligned FNC state vectors to EEG using the time point corresponding to the center of the sliding window. As that study was performed during awake conditions only (eyes open and eyes closed awake conditions), we did not have many EEG-based transitions within the subject wakeful state. Since the data from this study has a ground truth (EEG-based) hypnogram transition state vector informing of subject’s sleep stage, we performed a classification study using linear support vector machine (SVM) to see if the alignment of EEG hypnogram best corresponds to start or middle or in-between shift (3, 5 or 7 TRs) of dFNC state vector obtained from sliding window size of 30s. Prior to running SVM, the dFNC connectivity matrix (1891 pairs) was reduced to 30 dimensions using PCA. An 11 fold cross-validation was performed to assess the consistency of best alignment. For each fold, 5 different subjects were left out and a linear SVM was trained on remaining 50 subjects using libsvm package (https://www.csie.ntu.edu.tw/cjlin/libsvm/). A multi-class (all pairs/one-against-one) linear SVM model identifies support points in kernel space that maximally separates each pair of classes (W-N1,W-N2,W-N3,N1-N2,N1-N3,N2-N3). Then each test case/window is assigned to the class that gets most votes. Both training and test balanced (averaged per class) accuracies were computed. The two awake dFNC states were collapsed into one for this analysis.

To identify optimal window size among the tested window sizes, we performed another linear SVM separately for dFNC windows obtained with each choice of window length. Similar cross-validation analysis as mentioned above was used. Finally, we also tested the performance of the DCC algorithm, which uses adaptive windowing, to track subject sleep state. The DCC algorithm fits univariate generalized autoregressive conditional heteroscedastic (GARCH) GARCH(1,1) models to each univariate time series to obtain standardized residuals and then applies an exponentially weighted moving average window on these standard residuals of each pair of time courses to compute a non-normalized version of time-varying correlation between the two time courses (see (Lindquist et al., 2014; Choe et al., 2017) for more details).

To investigate if wakeful state can be further clustered into meaningful sub-clusters as in our earlier work, we restricted our clustering analysis to the 26001 dFNC windows (State 1 for window size 30) that corresponded to the subject wakeful EEG condition. This analysis revealed 4 sub-clusters with distinct connectivity profiles. We ran a SVM classification analysis by holding out 6000 dFNC windows and performed a three fold cross validation on the remaining 20000 windows to estimate a model across a range of parameter C [0.1 to 10] for a linear kernel. The best model was then used to test the classification accuracy on the held out samples and we computed average per class accuracy and confusion matrix.

### 2.6. Head motion effects

To assess the impact of subject head motion on the connectivity estimates, we assessed the relationship between the subject head movement summaries and dFNC state vector. We computed framewise displacement (FD) and framewise data variation (DVARS) (Power et al., 2012) for each subject to represent their motion summary. We then computed the number of instances subject DVARS exceeds its mean by 2.5 times its standard deviation for each k-means state. A one-way ANOVA was then performed on the counts to see if certain states show significantly more motion related outliers. In addition, we plotted each subject state vector along with their FD and DVARS to visually assess if any of the states are contaminated by head movement.

## 3. Results

The sixty-two ICNs selected for subsequent analysis are depicted in Figure 2. These components are grouped into subcortical (5), auditory (2), sensorimotor (10), visual (11), a set of higher order associative areas involved in attentional and executive control as well as cognitive control (19), default-mode regions (10) and cerebellar (5) components based on anatomical proximity and functional connectivity as in our earlier studies (Allen et al., 2012a,2011). The selected 62 ICN labels are summarized in Table 1.

**Figure 2:**
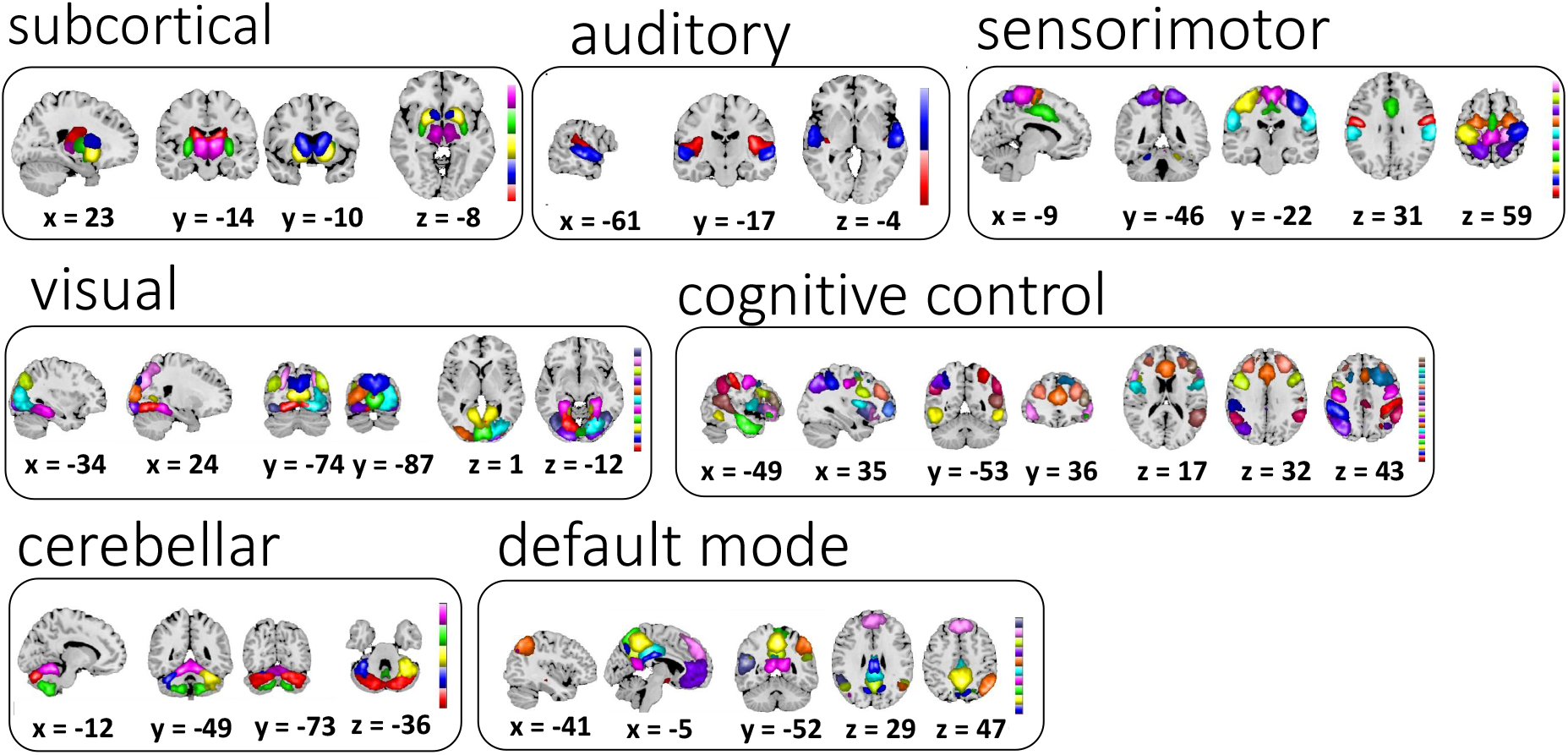
Sixty-two selected ICNs for further analysis were grouped into 7 modules using previously reported methods (Allen et al., 2012b).

**Table 1:**
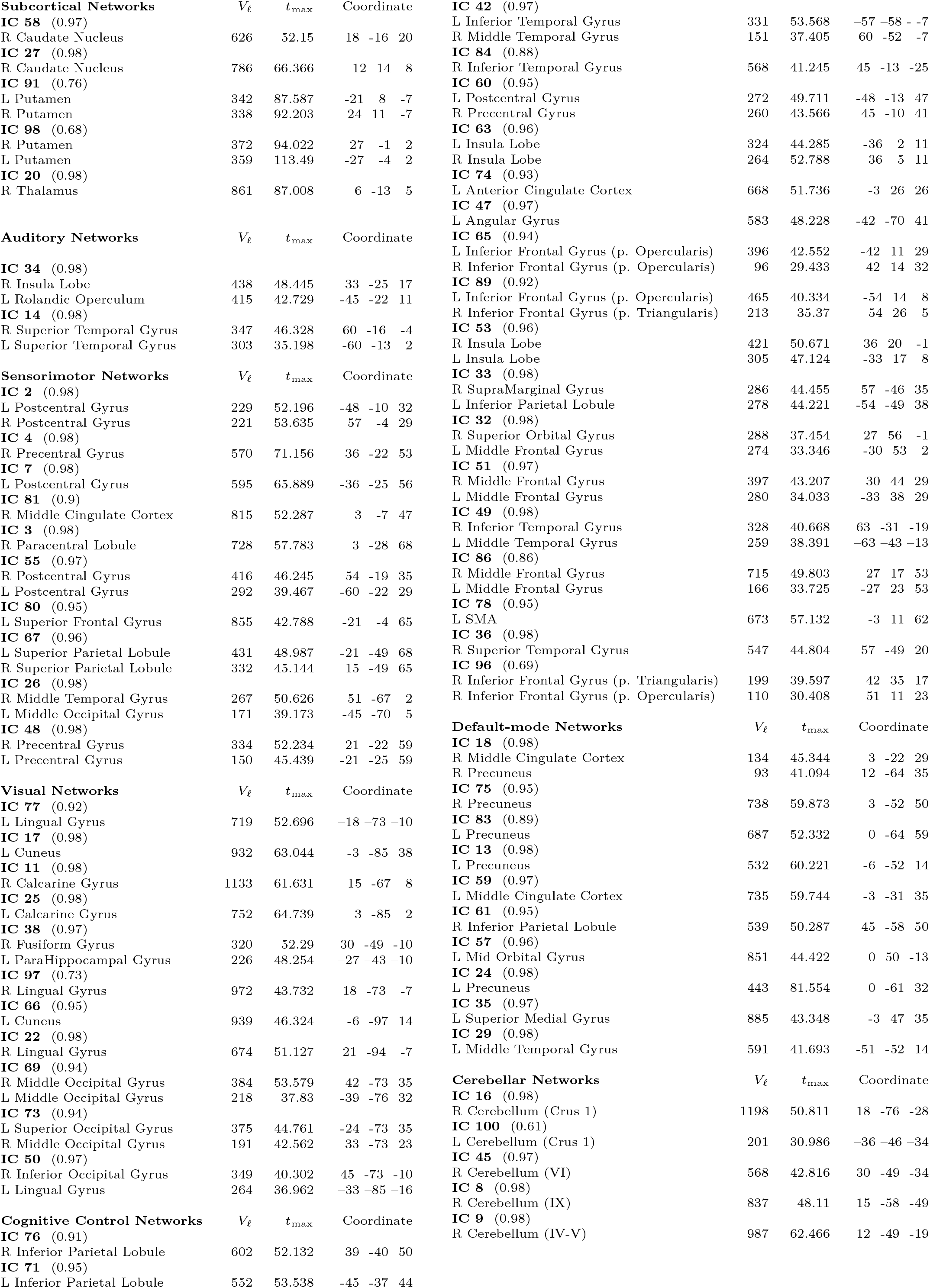
Peak activations of ICN SMs. The quality index (*I_q_*) associated with each ICN is listed in parentheses adjacent to the component number; *V_ℓ_* = number of vexels in each cluster; *t_max_* = maximum t-statistic in each cluster;Coordinate = coordinate (in mm) of *t_max_* in MNI space, following LPI convention.

|  |  |  |  |  |  |  |  |
| --- | --- | --- | --- | --- | --- | --- | --- |
| **Subcortical Networks** | $V_\ell$ | $t_{\max}$ | Coordinate | **IC 42** (0.97) | | | |
| **IC 58** (0.97) |  |  |  | L Inferior Temporal Gyrus | 331 | 53.568 | -57 -58 -7 |
| R Caudate Nucleus | 626 | 52.15 | 18 -16 20 | R Middle Temporal Gyrus | 151 | 37.405 | 60 -52 -7 |
| **IC 27** (0.98) |  |  |  | **IC 84** (0.88) |  |  |  |
| R Caudate Nucleus | 786 | 66.366 | 12 14 8 | R Inferior Temporal Gyrus | 568 | 41.245 | 45 -13 -25 |
| **IC 91** (0.76) |  |  |  | **IC 60** (0.95) |  |  |  |
| L Putamen | 342 | 87.587 | -21 8 -7 | L Postcentral Gyrus | 272 | 49.711 | -48 -13 47 |
| R Putamen | 338 | 92.203 | 24 11 -7 | R Precentral Gyrus | 260 | 43.566 | 45 -10 41 |
| **IC 98** (0.68) |  |  |  | **IC 63** (0.96) |  |  |  |
| R Putamen | 372 | 94.022 | 27 -1 2 | L Insula Lobe | 324 | 44.285 | -36 2 11 |
| L Putamen | 359 | 113.49 | -27 -4 2 | R Insula Lobe | 264 | 52.788 | 36 5 11 |
| **IC 20** (0.98) |  |  |  | **IC 74** (0.93) |  |  |  |
| R Thalamus | 861 | 87.008 | 6 -13 5 | L Anterior Cingulate Cortex | 668 | 51.736 | -3 26 26 |
|  |  |  |  | **IC 47** (0.97) |  |  |  |
| **Auditory Networks** | $V_\ell$ | $t_{\max}$ | Coordinate | L Angular Gyrus | 583 | 48.228 | -42 -70 41 |
|  |  |  |  | **IC 65** (0.94) |  |  |  |
| **IC 34** (0.98) |  |  |  | L Inferior Frontal Gyrus (p. Opercularis) | 396 | 42.552 | -42 11 29 |
| R Insula Lobe | 438 | 48.445 | 33 -25 17 | R Inferior Frontal Gyrus (p. Opercularis) | 96 | 29.433 | 42 14 32 |
| L Rolandic Operculum | 415 | 42.729 | -45 -22 11 | **IC 89** (0.92) |  |  |  |
| **IC 14** (0.98) |  |  |  | L Inferior Frontal Gyrus (p. Opercularis) | 465 | 40.334 | -54 14 8 |
| R Superior Temporal Gyrus | 347 | 46.328 | 60 -16 -4 | R Inferior Frontal Gyrus (p. Triangularis) | 213 | 35.37 | 54 26 5 |
| L Superior Temporal Gyrus | 303 | 35.198 | -60 -13 2 | **IC 53** (0.96) |  |  |  |
|  |  |  |  | R Insula Lobe | 421 | 50.671 | 36 20 -1 |
| **Sensorimotor Networks** | $V_\ell$ | $t_{\max}$ | Coordinate | L Insula Lobe | 305 | 47.124 | -33 17 8 |
| **IC 2** (0.98) |  |  |  | **IC 33** (0.98) |  |  |  |
| L Postcentral Gyrus | 229 | 52.196 | -48 -10 32 | R SupraMarginal Gyrus | 286 | 44.455 | 57 -46 35 |
| R Postcentral Gyrus | 221 | 53.635 | 57 -4 29 | L Inferior Parietal Lobule | 278 | 44.221 | -54 -49 38 |
| **IC 4** (0.98) |  |  |  | **IC 32** (0.98) |  |  |  |
| R Precentral Gyrus | 570 | 71.156 | 36 -22 53 | R Superior Orbital Gyrus | 288 | 37.454 | 27 56 -1 |
| **IC 7** (0.98) |  |  |  | L Middle Frontal Gyrus | 274 | 33.346 | -30 53 2 |
| L Postcentral Gyrus | 595 | 65.889 | -36 -25 56 | **IC 51** (0.97) |  |  |  |
| **IC 81** (0.9) |  |  |  | R Middle Frontal Gyrus | 397 | 43.207 | 30 44 29 |
| R Middle Cingulate Cortex | 815 | 52.287 | 3 -7 47 | L Middle Frontal Gyrus | 280 | 34.033 | -33 38 29 |
| **IC 3** (0.98) |  |  |  | **IC 49** (0.98) |  |  |  |
| R Paracentral Lobule | 728 | 57.783 | 3 -28 68 | R Inferior Temporal Gyrus | 328 | 40.668 | 63 -31 -19 |
| **IC 55** (0.97) |  |  |  | L Middle Temporal Gyrus | 259 | 38.391 | -63 -43 -13 |
| R Postcentral Gyrus | 416 | 46.245 | 54 -19 35 | **IC 86** (0.86) |  |  |  |
| L Postcentral Gyrus | 292 | 39.467 | -60 -22 29 | R Middle Frontal Gyrus | 715 | 49.803 | 27 17 53 |
| **IC 80** (0.95) |  |  |  | L Middle Frontal Gyrus | 166 | 33.725 | -27 23 53 |
| L Superior Frontal Gyrus | 855 | 42.788 | -21 -4 65 | **IC 78** (0.95) |  |  |  |
| **IC 67** (0.96) |  |  |  | L SMA | 673 | 57.132 | -3 11 62 |
| L Superior Parietal Lobule | 431 | 48.987 | -21 -49 68 | **IC 36** (0.98) |  |  |  |
| R Superior Parietal Lobule | 332 | 45.144 | 15 -49 65 | R Superior Temporal Gyrus | 547 | 44.804 | 57 -49 20 |
| **IC 26** (0.98) |  |  |  | **IC 96** (0.69) |  |  |  |
| R Middle Temporal Gyrus | 267 | 50.626 | 51 -67 2 | R Inferior Frontal Gyrus (p. Triangularis) | 199 | 39.597 | 42 35 17 |
| L Middle Occipital Gyrus | 171 | 39.173 | -45 -70 5 | R Inferior Frontal Gyrus (p. Opercularis) | 110 | 30.408 | 51 11 23 |
| **IC 48** (0.98) |  |  |  |  |  |  |  |
| R Precentral Gyrus | 334 | 52.234 | 21 -22 59 | **Default-mode Networks** | $V_\ell$ | $t_{\max}$ | Coordinate |
| L Precentral Gyrus | 150 | 45.439 | -21 -25 59 | **IC 18** (0.98) |  |  |  |
| **Visual Networks** | $V_\ell$ | $t_{\max}$ | Coordinate | R Middle Cingulate Cortex | 134 | 45.344 | 3 -22 29 |
| **IC 77** (0.92) |  |  |  | R Precuneus | 93 | 41.094 | 12 -64 35 |
| L Lingual Gyrus | 719 | 52.696 | -18 -73 -10 | **IC 75** (0.95) |  |  |  |
| **IC 17** (0.98) |  |  |  | R Precuneus | 738 | 59.873 | 3 -52 50 |
| L Cuneus | 932 | 63.044 | -3 -85 38 | **IC 83** (0.89) |  |  |  |
| **IC 11** (0.98) |  |  |  | L Precuneus | 687 | 52.332 | 0 -64 59 |
| R Calcarine Gyrus | 1133 | 61.631 | 15 -67 8 | **IC 13** (0.98) |  |  |  |
| **IC 25** (0.98) |  |  |  | L Precuneus | 532 | 60.221 | -6 -52 14 |
| L Calcarine Gyrus | 752 | 64.739 | 3 -85 2 | **IC 59** (0.97) |  |  |  |
| **IC 38** (0.97) |  |  |  | L Middle Cingulate Cortex | 735 | 59.744 | -3 -31 35 |
| R Fusiform Gyrus | 320 | 52.29 | 30 -49 -10 | **IC 61** (0.95) |  |  |  |
| L ParaHippocampal Gyrus | 226 | 48.254 | -27 -43 -10 | R Inferior Parietal Lobule | 539 | 50.287 | 45 -58 50 |
| **IC 97** (0.73) |  |  |  | **IC 57** (0.96) |  |  |  |
| R Lingual Gyrus | 972 | 43.732 | 18 -73 -7 | L Mid Orbital Gyrus | 851 | 44.422 | 0 50 -13 |
| **IC 66** (0.95) |  |  |  | **IC 24** (0.98) |  |  |  |
| L Cuneus | 939 | 46.324 | -6 -97 14 | L Precuneus | 443 | 81.554 | 0 -61 32 |
| **IC 22** (0.98) |  |  |  | **IC 35** (0.97) |  |  |  |
| R Lingual Gyrus | 674 | 51.127 | 21 -94 -7 | L Superior Medial Gyrus | 885 | 43.348 | -3 47 35 |
| **IC 69** (0.94) |  |  |  | **IC 29** (0.98) |  |  |  |
| R Middle Occipital Gyrus | 384 | 53.579 | 42 -73 35 | L Middle Temporal Gyrus | 591 | 41.693 | -51 -52 14 |
| L Middle Occipital Gyrus | 218 | 37.83 | -39 -76 32 |  |  |  |  |
| **IC 73** (0.94) | | | | **Cerebellar Networks** | $V_\ell$ | $t_{\max}$ | Coordinate |
| L Superior Occipital Gyrus | 375 | 44.761 | -24 -73 35 | **IC 16** (0.98) |  |  |  |
| R Middle Occipital Gyrus | 191 | 42.562 | 33 -73 23 | R Cerebellum (Crus 1) | 1198 | 50.811 | 18 -76 -28 |
| **IC 50** (0.97) |  |  |  | **IC 100** (0.61) |  |  |  |
| R Inferior Occipital Gyrus | 349 | 40.302 | 45 -73 -10 | L Cerebellum (Crus 1) | 201 | 30.986 | -36 -46 -34 |
| L Lingual Gyrus | 264 | 36.962 | -33 -85 -16 | **IC 45** (0.97) |  |  |  |
| **Cognitive Control Networks** | $V_\ell$ | $t_{\max}$ | Coordinate | R Cerebellum (VI) | 568 | 42.816 | 30 -49 -34 |
| **IC 76** (0.91) |  |  |  | **IC 8** (0.98) |  |  |  |
| R Inferior Parietal Lobule | 602 | 52.132 | 39 -40 50 | R Cerebellum (IX) | 837 | 48.11 | 15 -58 -49 |
| **IC 71** (0.95) |  |  |  | **IC 9** (0.98) |  |  |  |
| L Inferior Parietal Lobule | 552 | 53.538 | -45 -37 44 | R Cerebellum (IV-V) | 987 | 62.466 | 12 -49 -19 |

## 3.1. dFNC clustering results

The centroids (k=5) obtained from k-means clustering of dynamic FNC window data of all subjects are shown in Figure 3A. The centroids were ordered according to their frequency of occurrence in time (from the most awake state to the deepest sleep state), with the exception of state 2 Figure 3B. These centroids show distinct connectivity patterns from state to state. Estimation of modularity of the centroid states using the Louvain community detection algorithm (Rubinov and Sporns, 2011) resulted in three modules for states 1,2 and 3, and four modules for states 4 and 5. Although subcortical and cerebellar ICs belong to the same module in all states, they segregate as an additional module for states 4 and 5. We computed mean within module connectivity strength and the top 12 ICs for each module are shown in Figure 3C.

**Figure 3:**
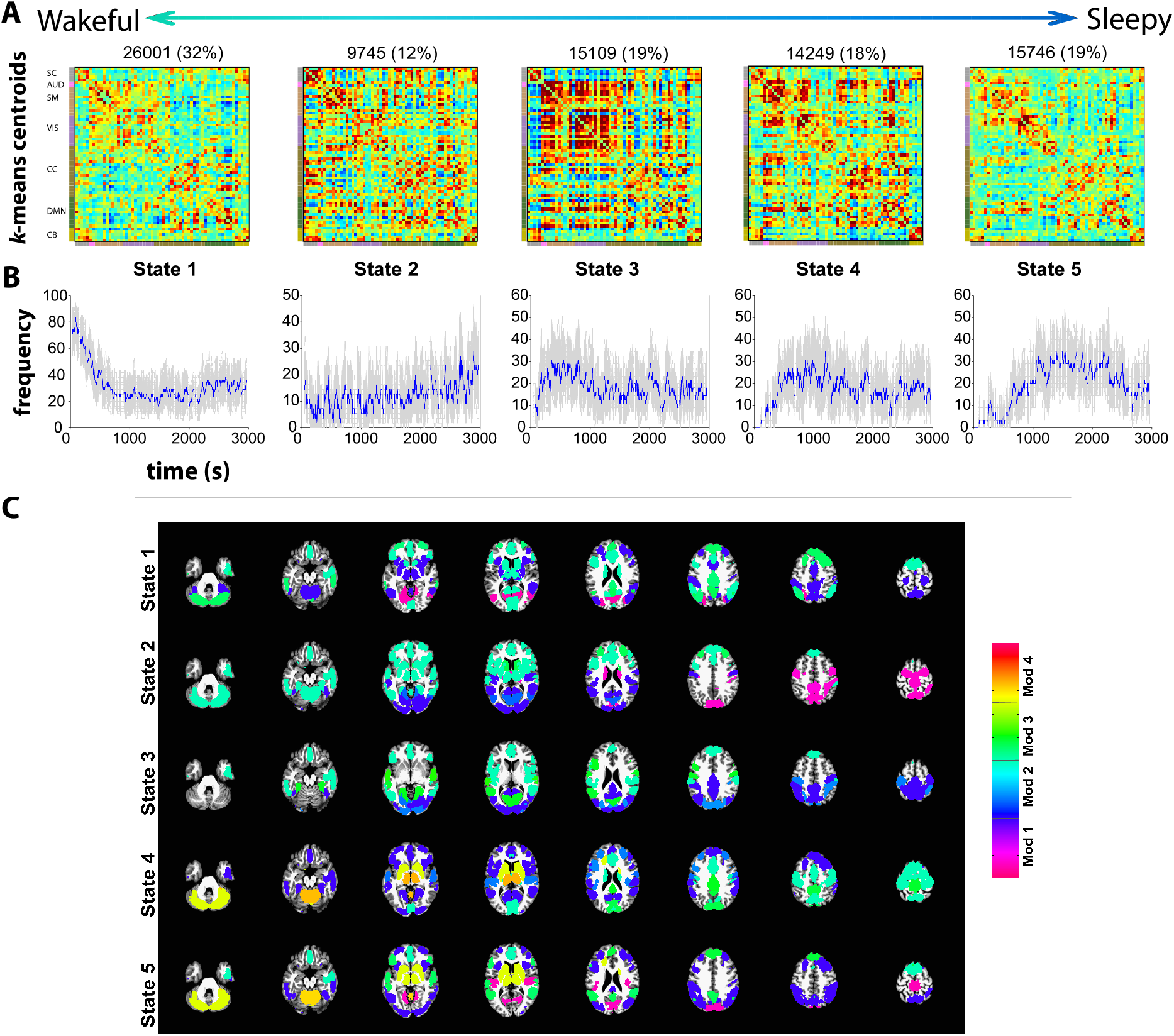
Cluster centroids from k-means clustering of dFNC window data for window size 30 (A) and the frequency of occurrence of each state in time (B). The standard errors for the frequency of occurrence were computed using 100 bootstrap resamples of the subject dFNC window data. (C) The modularity of the centroids is computed using the Louvain algorithm (“modularity_louvain_und_sign.m” function in the Brain Connectivity toolbox) resulting in three modules (Mod) for states 1,2 and 3 and four modules for states 4 and 5. The top 12 ICs with highest mean within module FC are depicted scaled by mean within module FC. Note that the weights (mean within module connectivity) are lower in states 1 and 5 compared to states 2, 3 and 4.

### 3.2. Do clustered connectivity states correspond to sleep states?

The subject-state vectors are sorted by sleep state (W, N1, N2 and N3) and frequency counts by sleep state were obtained and are shown in Figure 4. As seen in the figure, connectivity patterns in states 1 and 2 predominantly occur in the awake state while the patterns seen in states 3, 4 and 5 occur more frequently as subjects fall into different sleep states gradually with the connectivity pattern in state 5 occurring during N3 (deep) sleep stage.

**Figure 4:**
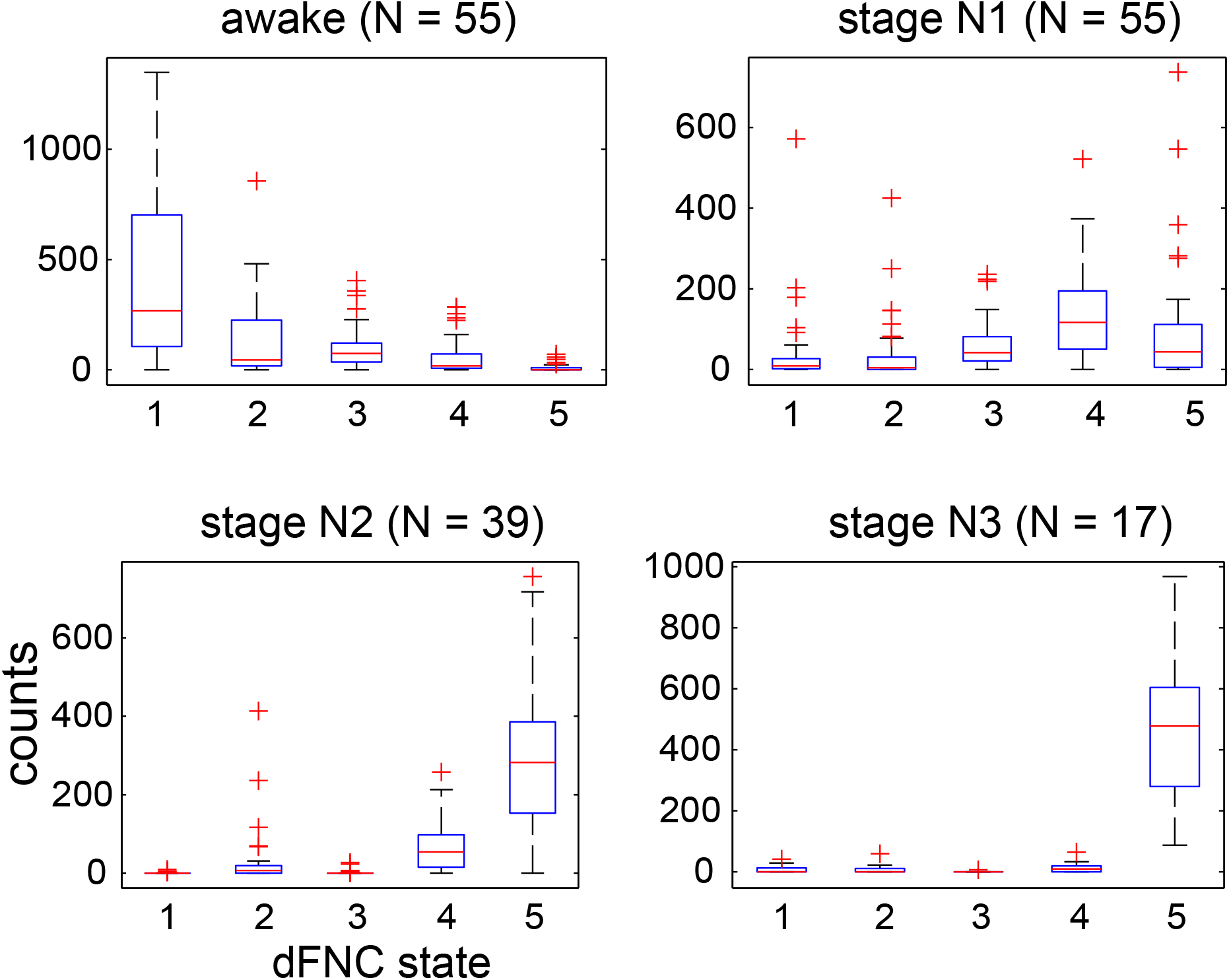
Frequency counts of state vector assignments obtained from k-means clustering of dFNC data sorted by hypnogram states. States 1 and 2 primarily occur during wakefulness, dFNC states 4 and 5 during N2 sleep stage and state 5 is predominant during deep sleep (N3 stage).

We tested the correspondence between subject state vectors obtained through clustering dFNC matrices and subject hypnograms obtained by computing the cross-correlation between the two vectors. Figure 5 shows examples of subjects with the two best and the two worst correlations between the two. As seen in the figure, the best subject showed a correlation of 0.89 between his/her hypnogram and state vector. The subjects that showed the poorest correlation between the two primarily tended to stay awake throughout the scan session, and the dFNC state vector showed transitions between awake related states 1 and 2.

**Figure 5:**
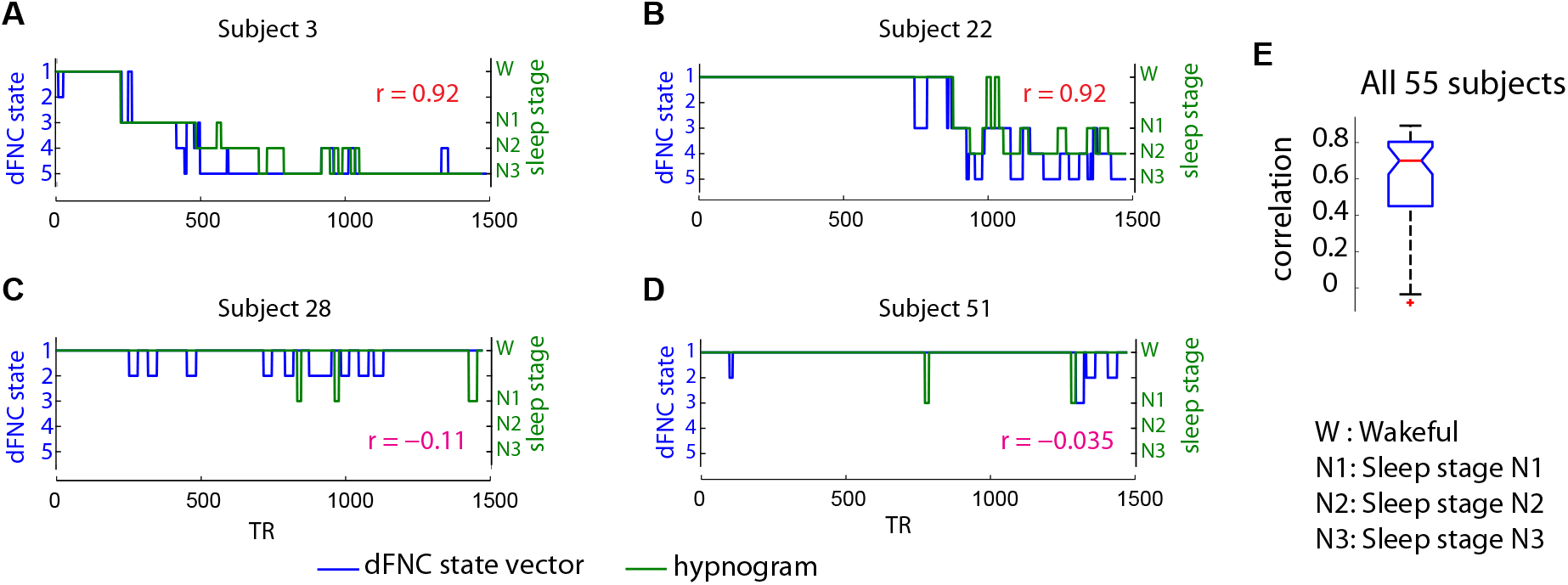
Comparison of subject state vectors obtained from k-means clustering of dFNC windows and EEG derived hypnograms for the two subjects with the highest correlation (A and B) and the two subjects with the lowest correlation (C and D). The overall distribution of the correlation between the two state vectors for all 55 subjects is presented in E. As seen, the subjects with low correlation tend to be awake throughout the scan session and the corresponding dFNC state vector transitions within the states 1 and 2 that are prevalent during wakefulness.

The results assessing the correspondence between subject dFNC state vector and his/her hypnogram are summarized in Figure 6. For this projection, a random sample of 400 dFNC windows by state (total 2000 points) were visualized using the t-SNE algorithm and were color coded with their corresponding k-means cluster assignment (Fig 6A) or by sleep stage obtained from the respective hypnogram (Fig 6B). Both awake states from the k-mean clustering are grouped together and show a transition from wakefulness to deeper sleep stages that occurs gradually along a smooth trajectory. This result shows that the dFNC estimates using the sliding window method and subsequent clustering correspond well to neurophysiological states as estimated via the EEG-based hypnogram.

**Figure 6:**
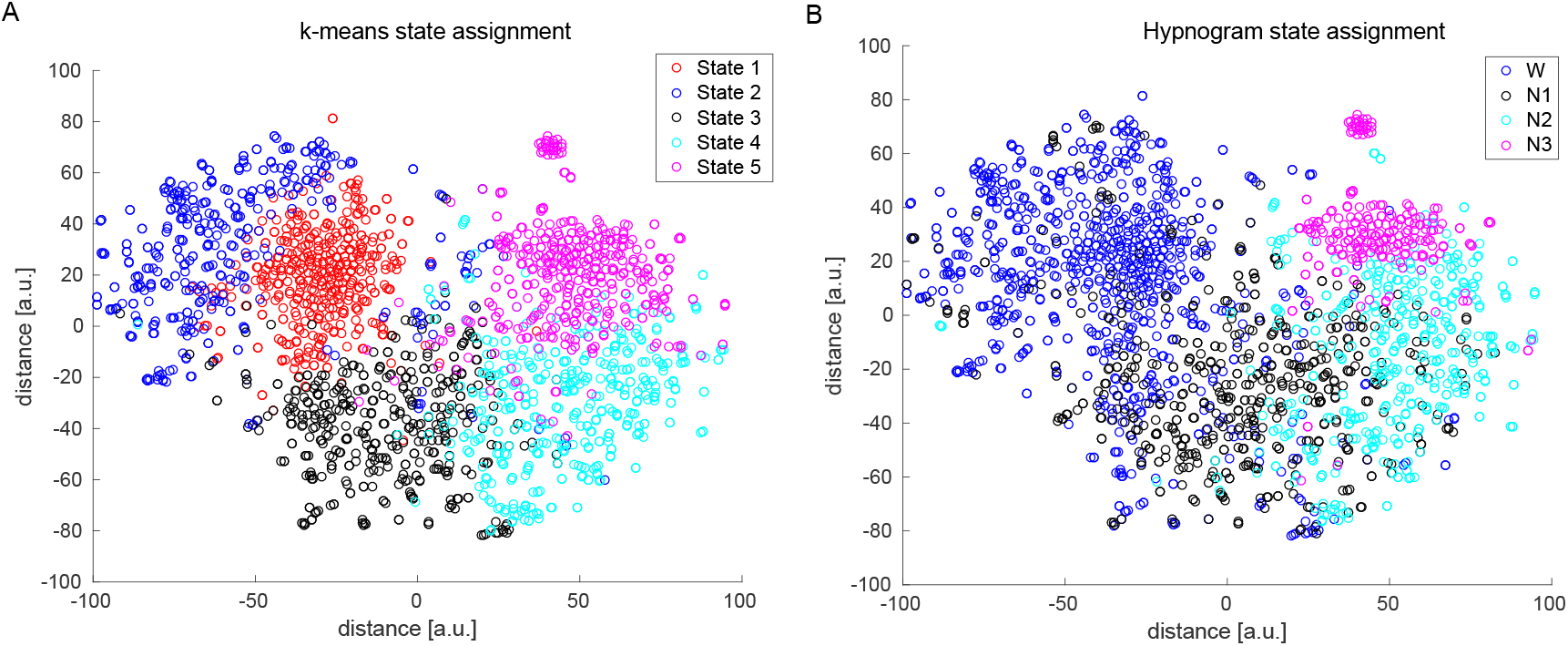
2D Visualization of dFNC data: We selected 2000 random dFNC windows (400 per dFNC state) and projected the multidimensional (1891) data into 2 dimensions using the t-SNE algorithm. The resulting mapping was subsequently color coded by k-means clustering assignment into 5 states (A) and by the subject EEG hypnogram state of that point (B). The data dimension was reduced to 30 principal components and a perplexity value of 35 was used for this projection.

### 3.3. How does motion affect the clustering?

To assess the effect of subject head movement during the scan on dFNC clustering results, we computed the number of windows with significant subject head movements (points greater that 2.5 standard deviations from mean framewise displacement) for each dFNC state and also visually assessed subject dFNC state vector and mean framewise displacement vectors. State 2 is associated with larger head movements during the scan relative to the other states. The number of significant head movements by state are shown in Supplemental Figure S1A. A couple of example subject state vectors and their head movement summary (FD) vectors are also shown in Supplemental Figure S1B.

### 3.4. Characterization of temporal dynamics

Comparison of the temporal properties of state vectors obtained from each modality is summarized in (Figure 7). The average dwell times and frequency of occurrence of each state are consistent for data from both modalities. The state transition matrices show good correspondence with more probable transitions from W->N1, N1->N2, N2->N3 and transitions to the W state from all sleep stages. This is in line with our knowledge of progression into various sleep stages. However, we observe a higher average number of transitions between states from the dFNC clustering derived state vectors compared to those computed using the EEG derived hypnogram.

**Figure 7:**
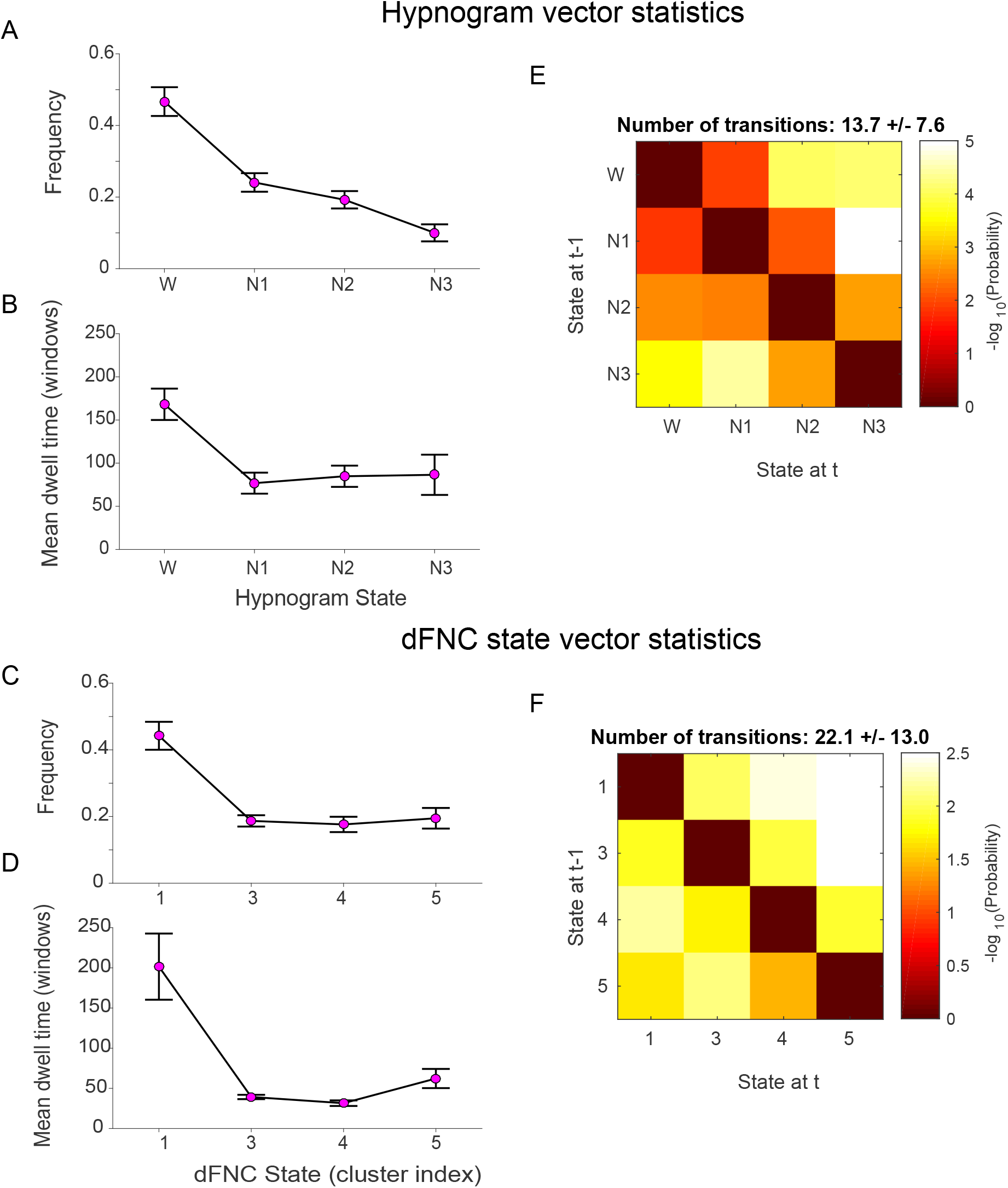
Comparison of state vector statistics and transition matrices computed from EEG-derived hypnograms and dFNC cluster derived subject state vectors. Hypnogram and dFNC state vector exhibit similar frequency of occurrences (A and C) and mean dwell times (C and D) respectively. The mean state transition matrices for hypnogram vectors (E) and dFNC state vectors (F) inform about the probability of transitioning from a given state *i* at time t-1 to state *j* at time t. The probabilities are converted to -log(10) scale, so higher (yellow to white) intensity values mean lower probability to transition. For both modalities, these matrices demonstrate tendency to remain in a given state (diagonal values are lower). The transitions to neighboring states are more likely in both the hypnogram and the dFNC state vectors. While there is chance of transitioning from deep sleep N3 to any other state (W, N1 or N2), the probability of transitioning from wakefulness at time t-1 immediately to deeper sleep stages (N2, N3) at time t is very low suggesting gradual transition from W to N3 stage. Note that dFNC states 1 and 2 are combined for this analysis.

### 3.5. Alignment of EEG derived hypnogram and windowed dFNC data

The SVM classification results for best alignment of EEG hypnogram and dFNC window position are presented in Figure 8. The results suggest that both start and middle TR of dFNC window aligned to corresponding EEG hypnogram vector results in similar classification performance. However in-between shifts resulted in poor classification accuracy.

**Figure 8:**
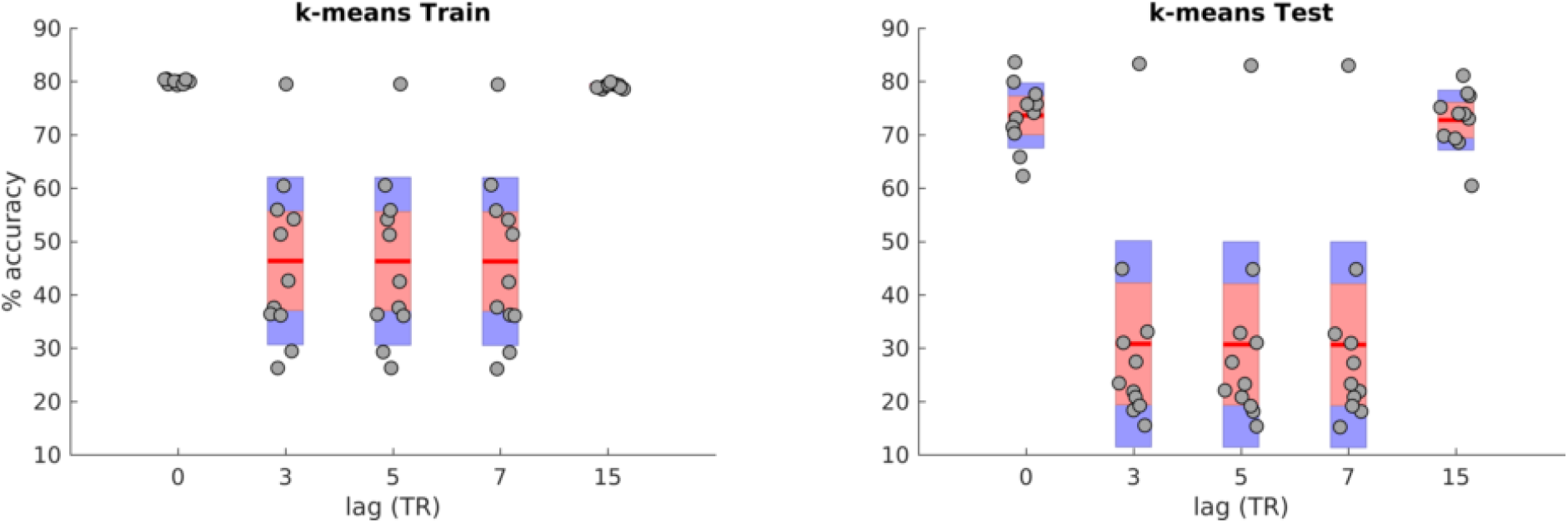
SVM classification accuracies for training (A) and test (B) cases of alignment between EEG hypnogram and subject dFNC state vector obtained using a window size of 30 TRs. The alignment is tested for lags 0, 3, 5, 7 and 15 TRs. Each point in the distribution corresponds to the balanced per class accuracy from one of the 11 cross-validation iterations of training data (left) that included data from 50 random subjects and the accuracy from left of test data (right) that included data from 5 remaining subjects.

### 3.6. How well can we predict sleep stages from dFNC data?

The SVM classification accuracies comparing the prediction of subject hypnogram with dFNC estimates obtained using different window lengths is presented in Figure 9. As seen in the figure, the classification accuracy significantly increases with dFNC estimates from shorter to longer window sizes in training subject cases (the data SVM model has seen: one-way ANOVA F=342,p<1e-35). The accuracies on left out test samples did not significantly differ with window sizes (one-way ANOVA F=0.7,p>0.58). These accuracy rates are consistent with earlier reports (Tagliazucchi et al., 2012b) however we show that these can be achieved using much shorter window lengths. The classification accuracies for dFNC estimates using DCC method performed poorly compared to those obtained using sliding window methods for all window sizes.

**Figure 9:**
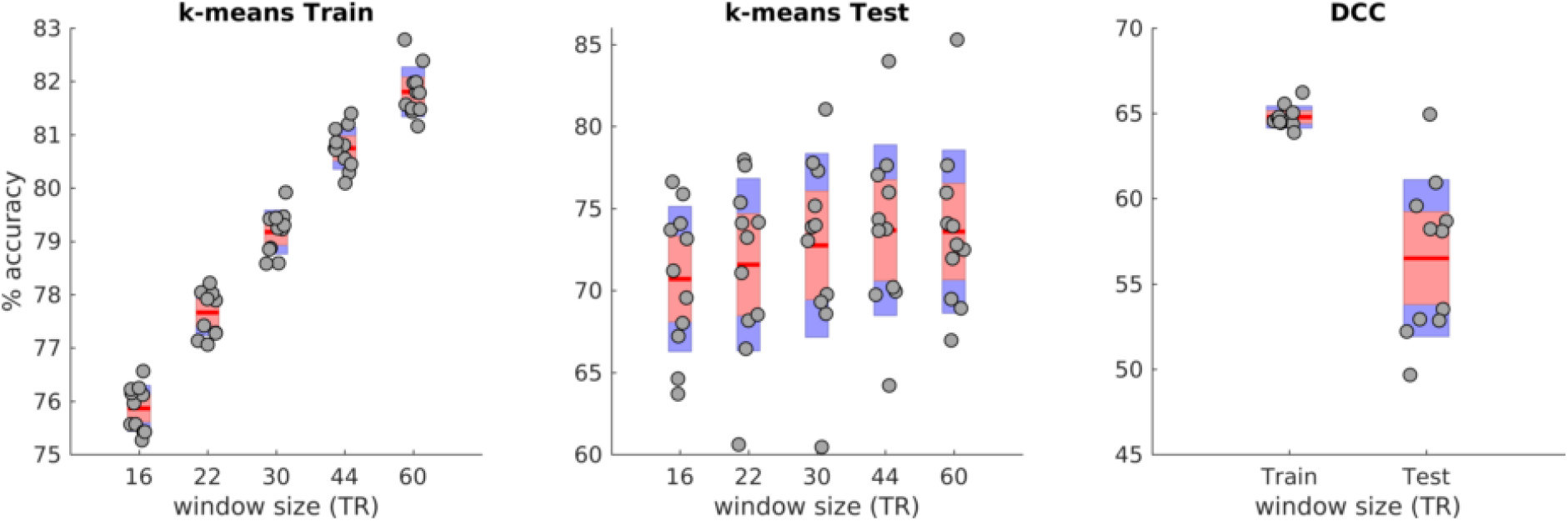
Linear SVM classification accuracies of subject sleep stage from dFNC estimates obtained using different window sizes for training (left), test (middle) data from 11 cross-validation iterations and the classification accuracies obtained from the DCC estimates for the same cross-validation scheme for training and left out test data are presented in the right.

### 3.7. Does wakeful stage correspond to only one dFNC cluster?

Since our prior work (Allen et al., 2017) showed multiple wakeful states with distinct EEG spectral signatures, we further focused on the awake condition only to see if it can be reliably segmented into sub-clusters. A search for the optimal number of clusters using the elbow criterion yielded four clusters. The cluster centroids are depicted in Figure 10. Awake state cluster centroids 1 and 2 resemble each other but differ in the strength of correlations in within and between module groupings. Awake state 4 resembles state 3 from the full dataset but distinguishes itself in anti-correlations between sensory (visual, motor and auditory) networks to higher order cognitive networks and also to the default-mode regions. Results from SVM classification of awake only cluster windows resulted in 92% classification accuracy using a one-vs-rest linear SVM model with parameter C=1. The resulting confusion matrix is presented in Table 2. The classification accuracy reaches chance level of 25% when the class labels are permuted. This result suggests that the sub-clusters obtained from awake state are linearly separable with high accuracy.

**Figure 10:**
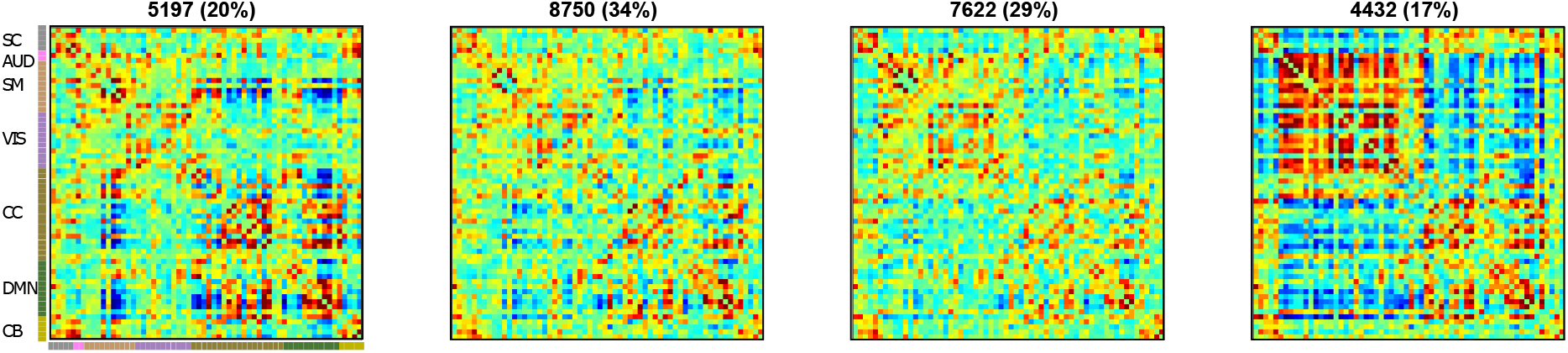
Cluster centroids from k-means clustering of window dFNC data of awake only state (State 1). The estimated clusters were observed to have meaningful and distinct structure.

**Table 2:** Awake state SVM classification confusion matrix. The percent classification accuracy from linear SVM model for each of the four awake only K-means states. The high accuracy strongly suggests that these clusters are unlikely to be noise.

|  |  | Predicted |  |  |  |
| --- | --- | --- | --- | --- | --- |
|  |  | AwState 1 | AwState 2 | AwState 3 | AwState 4 |
| Actual | AwState 1 | 91.8 | 4.2 | 3.4 | 0.6 |
|  | AwState 2 | 0.6 | 94.8 | 3.4 | 1.2 |
|  | AwState 3 | 1.9 | 5.5 | 89.7 | 2.7 |
|  | AwState 4 | 0.9 | 2.8 | 3.8 | 92.4 |

## 4. Discussion

In this work, using an ICA based pipeline, we assess the ability of sliding window correlation based dynamic functional network connectivity measures to capture neurophysiological state transitions obtained from sleep staging of EEG data that was concurrently acquired during resting fMRI acquisition. Results show a good correspondence between the subject state vectors obtained from k-means clustering of dFNC windows and subject hypnograms. We further demonstrate that distinct resting functional connectivity patterns are associated with wakeful and sleep states with dFNC state 1 predominantly occurring while subjects are awake, dFNC state 3 corresponding to reduced subject vigilance and early sleep stage (N1) and dFNC states 4 and 5 are more likely to be associated with deeper sleep stages. Deep sleep (N3) is primarily associated with dFNC state 5 across all subjects. One state (dFNC state 2) primarily captures variance associated with subject movement.

### 4.1. dFNC clustering estimates can reliably predict subject sleep state

Recently, using the same data reported in this work, Haimovici et al. (2017) showed good correspondence between centroids of windowed resting fMRI correlation data obtained from k-means clustering and those informed by an EEG-based hypnogram. In Haimovici et al. (2017), the connectivity measures were estimated from non-overlapping windows of length 100 seconds within fixed regions of interest and so precludes time varying connectivity information. Our work replicated the aforementioned result in Haimovici et al. (2017) despite several differences including the use of time-varying connectivity estimates from ICA derived network time courses as well as the use of overlapping windows with a step size of 1 TR. Using this approach, we show that high classification accuracies can be obtained for windows as short as 30 seconds (15 TRs).

Our results also demonstrated that dFNC estimates from sliding window correlation show higher accuracy compared to an adaptive windowless dynamic conditional correlation method. However, DCC estimates from sliding window based covariance estimates resulted in classification accuracies similar to those obtained from the sliding window method suggesting this parametric model of conditional correlation for resting fMRI data might be more sensitive to noise than smoother sliding window estimates when instantaneous conditional correlation estimates are computed.

Our analysis demonstrates that even window lengths as short as 30 seconds result in reasonable estimates of time-varying connectivity profiles with estimated dFNC clustering states showing good classification accuracy with subject hypnogram. This is in line with previous reports (Shirer et al., 2012) and suggests that the 1/*f_min_* recommendation for window length as recommended by Leonardiand Van De Ville (2015) might be too conservative (Vergara and Calhoun, 2018). In our case 1/*f_min_* would result in 100 sec (since *f_min_* = 0.01 Hz after filtering) and may limit our ability to capture dynamics for 1/*f* spectral distributed BOLD data (Zalesky and Breakspear, 2015).

The observed differences between the individual dFNC state vector and the hypnogram can arise for the following reasons. The hypnogram is scored from EEG data epochs of 30 seconds and assigns each 30 second segment to a single hypnogram stage (W/N1/N2 or N3). The dFNC at a given instant is estimated from a window length ‘w’ on either side of the time point (past and future). However, k-means is a hard clustering approach and, as seen in Figure 6, there is ambiguity in the assignment of the data point with its immediate neighboring state (for example dFNC states 4 and 5 and corresponding N2 and N3 sleep stages) compared to the ground truth (hypnogram). Fuzzy k-means approaches can help mitigate this issue by allowing for states to overlap with one another (Miller et al., 2016). Another source of possible differences may be due to the different impact of noise (e.g. motion, MRI gradients) on the two signals which can hinder our ability of accurately estimating subject states.

### 4.2. Head motion appears to be separable from dynamic connectivity measures

Our dFNC clustering analysis reveals that motion appears to be mostly associated with one of the states (dFNC state 2) rather than spread across all of them. An examination of classification errors by sleep stage obtained for a window length of 30 TRs suggested that the awake state has the least errors (about 10%) and N1 sleep stage had the most errors (approximately 40%), with deeper sleep stages having an error rate of about 20%. Further evaluation to determine if dFNC windows overlapping with larger head movements during the scan drive these errors suggested that motion only contributed to about 15% of misclassified cases evenly across wakeful and sleep stages i.e 15% of 10, 40, 20 and 20 error rates of W, N1, N2, and N3 stages respectively. This suggests that dFNC windows exhibit larger variability during the N1 sleep stage compared to other sleep stages as observed in Supplemental Figure S2 leading to misclassification. This highlights the need to further identify additional sub-clusters in the N1 sleep stage and investigate if a fine grained EEG classification of this stage as proposed by Hori and colleagues (Hori et al., 1994) and recently demonstrated using EEG data (Jagannathan et al., 2018) can provide additional insights into large scale connectivity changes during the transition to sleep (Goupil and Bekinschtein, 2012).

### 4.3. Evidence of multiple dFNC sub-clusters states during wakefulness

The elbow criterion, computed as a ratio of between-cluster to within-cluster distance, was used to obtain an optimal k for clustering of the data using the k-means algorithm. Results suggest a 5 cluster solution within which we observe fewer (two) states from the wakeful portions of the scans than those reported earlier for data collected during the wakeful condition only (Allen et al., 2012b, 2017). This could be due to the fact that variability in dFNC fluctuations during wakefulness is lower compared to variability across different sleep stages (see Supplemental Figure S2). To evaluate this further, we performed a separate elbow criterion search and clustering of dFNC windows within only the wakeful state to see if additional clusters of meaningful time-varying connectivity profiles within and across subjects can be reliably estimated. Separate K-means clustering of dFNC windows from the awake only state (state 1) revealed additional sub-clusters not seen from clustering the data including all sleep stages. Windows corresponding to these sub-clusters are linearly separable with good accuracy using a linear SVM classifier. This result is in line with and extends recent reports showing replicable dFNC states in multiple independent datasets (Abrol et al., 2017) during (unconfirmed) wakeful conditions.

Some recent studies have argued that the observed connectivity states during wakefulness are primarily a reflection of sampling variability, changes in subject vigilance and partly reflect changes due to head movement (Laumann et al., 2016; Liegeois et al., 2017; Haimovici et al., 2017). Spontaneous eye blinks during wakefulness have also been shown to cause connectivity fluctuations in resting fMRI data (Wang et al., 2016). Another view is that time-varying connectivity changes in resting state fMRI can be modeled as hierarchical transitions between connectivity states consisting of two metastates: one corresponding to states with increased connectivity in brain regions involved in higher cognition and other corresponding to states with greater integration within sensory regions (Vidaurre et al., 2017). Another recent study identified strong correlation between dFC obtained using sliding window correlation of BOLD data and temporal dynamics of calcium signal during rest in mice brain recordings suggesting a neuronal origin of the observed dynamics (Matsui et al., 2018). In this work, we show distinct connectivity states during wakefulness that are separable via a cross-validated linear classifier. Future studies should evaluate the underlying neurobiological signatures of both sleep and wakefulness in greater detail.

### 4.4. Limitations and future work

We did not compare the sliding window correlation method to alternative methods of dynamic connectivity methods like multiplication of temporal derivatives (Shine et al., 2015) and time-frequency approaches (Yaesoubi et al., 2015b). For classification analysis, we used linear SVM with cross-validation. The classification accuracies might improve with a non-linear (RBF or polynomial) kernel with additional optimization of the kernel parameter search space. Also, we reduced the functional connectivity matrix dimension using PCA before training the SVM model. Alternative feature selection models like recursive feature elimination might improve classification accuracy. However we expect that the observed differences in accuracy of classification from dFNC data for different window lengths would remain even after employing more complex procedures. Also, in this work we show the presence of linearly separable (predictable) connectivity states during wakeful rest to extend our previous work and address some current technical controversies in the field. Going forward, it would be interesting to perform a hierarchical analysis of various sleep stages to evaluate the possibility of different states existing during different sleep stages as well as to further study the underlying neurobiological correlates of these states with the use of multimodal imaging data along with novel modeling techniques.

### 4.5. Conclusions

In this work, using an ICA-based pipeline applied to concurrent EEG/fMRI data collected during wakefulness and various sleep stages we demonstrate that time varying connectivity estimates from sliding windowed correlations of resting state functional network time courses well classify the sleep states obtained from EEG data even for windows as short as 30 seconds. We show that head motion is mostly associated with one of the states rather than spread across all of them. Consistent with earlier work, we find increased variability in connectivity as subjects transition from wakefulness to sleep. We report linearly separable clusters within the wakeful state and suggest future directions for assessing their neurobiological relevance via hierarchical analysis of predictable states in various sleep stages measured with EEG-fMRI data including eye tracking during the wakeful condition.

## 4.6. Acknowledgements

This work was funded by NIH P20GM103472, Ro1EB006841 and NSF EPSCoR 1539067 and the Bundesministerium für Bildung und Forschung (grant 01 EV 0703) and the LOEWE Neuronale Koordination Forschungsschwerpunkt Frankfurt (NeFF).

## 5. Supplementary data

**Figure S1:**
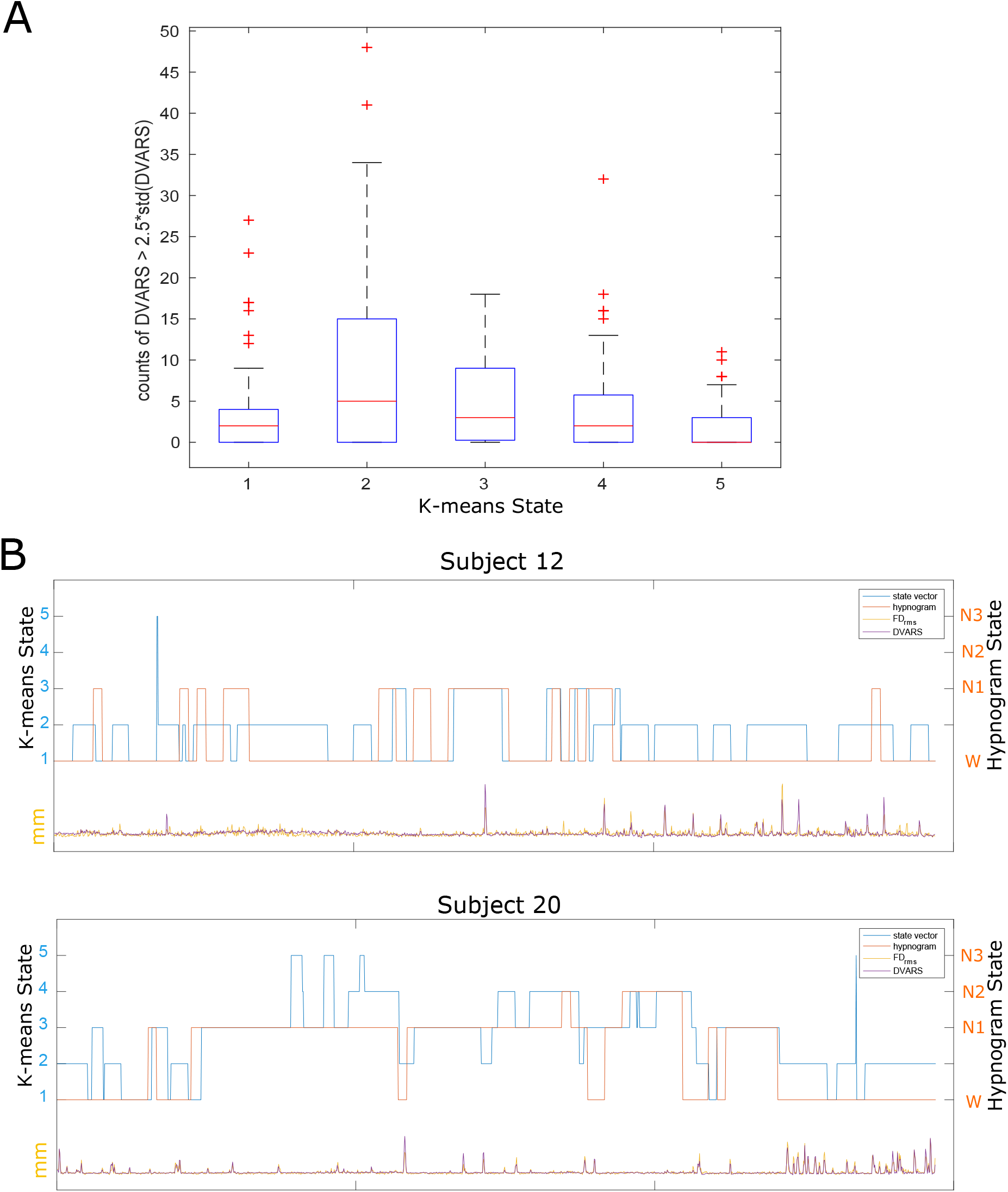
A) Count of number of DVARS of raw data exceeding 2.5 times its standard deviation by K-means state assignment. One-way ANOVA on mean differences in counts is significant with F=8.9 and *p* < 1e-05. A similar result is observed using subject FD values. B) Example subject state vector plotted along with his/her hynogram and head movement summaries (DVARS and FD). As seen, State 2 FC estimates seems to be contaminated by bigger jerky movements from subjects.

**Figure S2:**
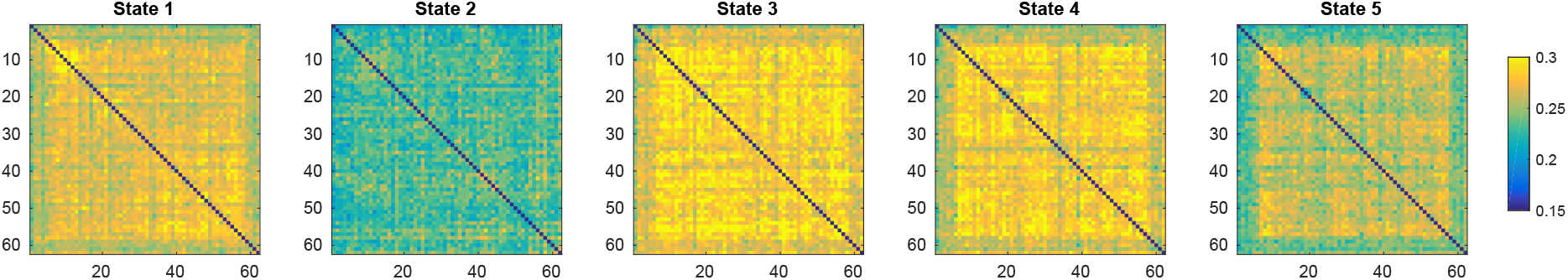
Standard deviation of the dFNC estimates by K-means state. The standard deviation increases for most pairs for States 3 and 4 compared wakeful state 1 and the variability of dFC reduces during deep sleep stage N3.

